# ATG7 vs. ATG5: Distinct Autophagy Pathways Shaping TGF-β Signaling and Endothelial Function

**DOI:** 10.1101/2025.10.15.682686

**Authors:** Aman Singh, Bharatsinai Peddi, Kriti S. Bhatt, Hien C. Nguyen, Krishna K. Singh

## Abstract

Autophagy and transforming growth factor-beta (TGF-β) signalling are critical cellular processes that maintain homeostasis and regulate various physiological functions, including endothelial cell function. This study explores the distinct roles of two essential autophagy regulators, autophagy-related gene 5 (ATG5) and ATG7, in modulating TGF-β signalling and endothelial function. To this end, ATG5 and ATG7 were selectively silenced in endothelial cells, with knockdown efficiency confirmed via RT-qPCR and Western blot analyses. Gene expression profiling revealed differential regulation of TGF-β signalling components following ATG5 or ATG7 silencing. Functional assays demonstrated that ATG5 knockdown enhanced endothelial cell proliferation and migration, whereas ATG7 silencing produced distinct, less pronounced effects. Furthermore, downstream effectors of the TGF-β pathway exhibited gene-specific modulation, underscoring divergent roles of ATG5 and ATG7 in this signalling cascade. Collectively, these findings highlight the non-redundant functions of ATG5 and ATG7 in coordinating TGF-β signalling pathways, offering new insights into their contribution to endothelial physiology and potential as therapeutic targets in vascular pathologies.

## Introduction

Autophagy is a catabolic cellular process responsible for the degradation and recycling of intracellular components such as damaged organelles and misfolded proteins via lysosomes and late endosomes [1]. A key feature of this process is the formation of a double-membraned structure called the phagophore, which typically originates at the surface of the endoplasmic reticulum. The phagophore elongates by incorporating lipids and autophagy-related proteins (ATGs), ultimately sealing to form the autophagosome, which engulfs cytoplasmic cargo for degradation. Two essential components of the core autophagy machinery are ATG5 and ATG7, which play distinct but cooperative roles in autophagosome formation [2]. ATG7 acts as an E1-like activating enzyme in two ubiquitin-like conjugation systems: the ATG12–ATG5 pathway and the LC3 (ATG8) lipidation pathway [3, 4]. In this cascade, ATG7 first activates ATG12, which is transferred to the E2-like enzyme ATG10 and conjugated to ATG5. The ATG12–ATG5 conjugate then associates with ATG16L1 to form a complex that functions as an E3-like ligase, facilitating the lipidation of LC3 with phosphatidylethanolamine (PE), producing LC3-II [5, 6]. LC3-II is critical for autophagosome membrane expansion, cargo recognition, and maturation. In the absence of ATG7, neither the ATG12–ATG5 complex nor LC3-II is generated, leading to a complete block in autophagosome biogenesis [7]. Similarly, deletion of ATG5 disrupts ATG12 conjugation and downstream LC3 lipidation, also resulting in loss of autophagosome formation [3]. Thus, ATG5 and ATG7 are indispensable for the initiation and progression of canonical autophagy, and the loss of either protein leads to a complete disruption of the autophagic process [8].

Impaired autophagy has emerged as a critical contributor to the development and progression of cardiovascular diseases (CVDs), particularly through its effects on endothelial cell function. Endothelial cells line the interior of blood vessels and are key regulators of vascular tone, inflammation, and barrier integrity. Autophagy in endothelial cells is essential for maintaining cellular homeostasis by removing damaged organelles, mitigating oxidative stress, and modulating inflammatory responses. When autophagy is disrupted, endothelial cells accumulate dysfunctional mitochondria and reactive oxygen species (ROS), leading to endothelial dysfunction—a hallmark of early CVDs [9, 10]. Impaired endothelial autophagy has been linked to increased vascular permeability, elevated expression of adhesion molecules, and a shift toward a pro-inflammatory and pro-thrombotic phenotype, all of which contribute to vascular inflammation and atherogenesis [11, 12]. Moreover, genetic deletion of key autophagy genes, such as ATG5 or ATG7 in endothelial cells exacerbates vascular injury, impairs angiogenesis, and accelerates atherosclerotic lesion formation in experimental models [13, 14]. These findings underscore the protective role of endothelial autophagy in vascular health and highlight its potential as a therapeutic target in cardiovascular pathology.

We previously demonstrated that endothelial cell-specific deletion of ATG7 results in a complete loss of autophagic activity in endothelial cells, which in turn activates the transforming growth factor-beta (TGF-β) signalling pathway and promotes endothelial-to-mesenchymal transition (EndMT) [8]. EndMT is characterized by the downregulation of endothelial markers (e.g., VE-cadherin and CD31) and upregulation of mesenchymal markers such as α-SMA and fibronectin, contributing to fibrosis and vascular pathology. Interestingly, although both ATG5 and ATG7 are essential for autophagosome formation and their deletion leads to a loss of canonical autophagy, we observed that only ATG7 loss induces EndMT, despite similar increases in the expression of TGF-β1 and its receptors TGFBR1 and TGFBR2. This divergence suggests that ATG5 and ATG7 may differentially regulate TGF-β signalling and endothelial plasticity through autophagy-dependent and possibly autophagy-independent mechanisms. While ATG7 appears to act as a critical gatekeeper of TGF-β-induced EndMT, ATG5 may play a more modulatory role, not sufficient on its own to trigger mesenchymal transition. These findings underscore the non-redundant roles of ATG proteins in endothelial biology and warrant further mechanistic studies to delineate the distinct pathways through which ATG5 and ATG7 influence endothelial function and vascular remodeling.

## Methods

### Cell Culture

Human Umbilical Vein Endothelial Cells (HUVECs, Lonza, mixed, passage 4-6) were cultured in an endothelial cell growth medium (EGM™-2 Bulletkit™; Lonza) supplemented with growth factors, 5% fetal bovine serum (FBS), and antibiotics at 37^°^C and 5% CO2. HUVECs were transfected with siATG7, siATG5 or scrambled control using the Lipofectamine™ 3000 transfection reagent (Invitrogen) according to the manufacturer’s protocol as previously described [8]. Briefly, 2×10^5^ HUVECs were seeded in 6-well cell culture plates and transfected with 5 nM siATG7, siATG5 or scrambled control (n=4/group) and incubated in endothelial cell growth medium. RNA and proteins were collected 24- and 48-hours post-transfection, respectively [8].

### Western Blot Analysis

Cell lysates were collected and extracted in RIPA buffer 48 hours post-transfection with siATG5, siATG7, or a scrambled control. Bradford assay was performed to calculate the concentration of the protein samples. SDS-polyacrylamide gels were run by loading equal amounts of proteins, which were then transferred to the PVDF membranes. The membranes were blocked for 1 hour in 5% BSA (Bovine Serum Albumin) and then incubated with primary antibodies for 2 hours at room temperature or at 4°C overnight. The primary antibodies used for the experiments include ATG5 (Cell Signalling, Cat #12994), ATG7 (Cell Signalling, Cat #8558), p21 (Cell Signaling # 2947S), SMAD3 (Cell Signalling, Cat #9523), and GAPDH (Cell Signaling # 2118S). The proteins were then incubated with secondary antibody for 1 hour at room temperature. Bands were visualized with the ECL substrates using a chemiluminescence channel and 700 channels in the LiCor Fc Odyssey system. The bands were quantified using LiCor Fc Odyssey inbuilt software.

### cDNA synthesis and Real-Time Quantitative PCR (RT-qPCR)

First, total RNA was collected in Trizol Reagent from the HUVECs following standard protocols. RNA concentration was measured using NanoDrop. Subsequently, cDNA synthesis was performed using Quantitect cDNA synthesis kit (Qiagen). Then, SYBR® Green–based RT-qPCR was conducted using the forward and reverse primers of *ATG5, ATG7, p21, TGFB1*, and *GAPDH* according to manufacturer’s instructions[8].

### PCR Array

Quantitative real-time PCR analysis of 84 TGF-β-related genes was performed with the human TGF-β /BMP signalling pathway RT^2^ profiler PCR array (PAHS-035, Qiagen). Data were analyzed with the manufacturer’s integrated web-based software package.

### Cell Proliferation

To measure endothelial cell proliferation of siATG5-, siATG7-, and scrambled control-transfected HUVECs, a WST-1 assay was performed. WST-1 is a red tetrazolium salt which is cleaved by metabolically active cells to formazan, which is dark red in colour; by quantifying the amount of formazan formed, we can measure the number of proliferating cells. HUVECs from the aforementioned conditions were seeded into 96-well plates for 48 hours, and 10μL of WST-1 reagent was added to each well. Cells were incubated at 37°C with 5% CO2 for 4 hrs. After incubation, the absorbance of the formazan was measured at 450 nm. A reference wavelength of 655 nm was used.

### Migration Assay

Cultured HUVECs were transfected and seeded at a density of 1.2 × 105 cells/well in a 6-well plate for 48 h. Each well was then administered a consistent straight scratch using sterilized ruler and a 200-μl tip prior to high-or normal glucose exposure. Phase contrast microscopy using an adapted camera (Optika) was employed to take images of scratch area at T0 and every 4 hrs up to 20 hrs. The wound healing tool of ImageJ was used for quantification of scratch area and calculation of migration rate as described [15].

## Results

### Validation of ATG7 and ATG5 Silencing at Transcript and Protein Levels

To confirm effective knockdown of ATG7 and ATG5, both RT-qPCR and Western blot analyses were performed to assess mRNA and protein expression levels, respectively. As shown in Figure 1A and 1B, ATG7 silencing was successfully achieved at both the transcript and protein levels. Similarly, Figure 1C and 1D demonstrate efficient knockdown of ATG5 at the corresponding molecular levels.

**Figure 1.**
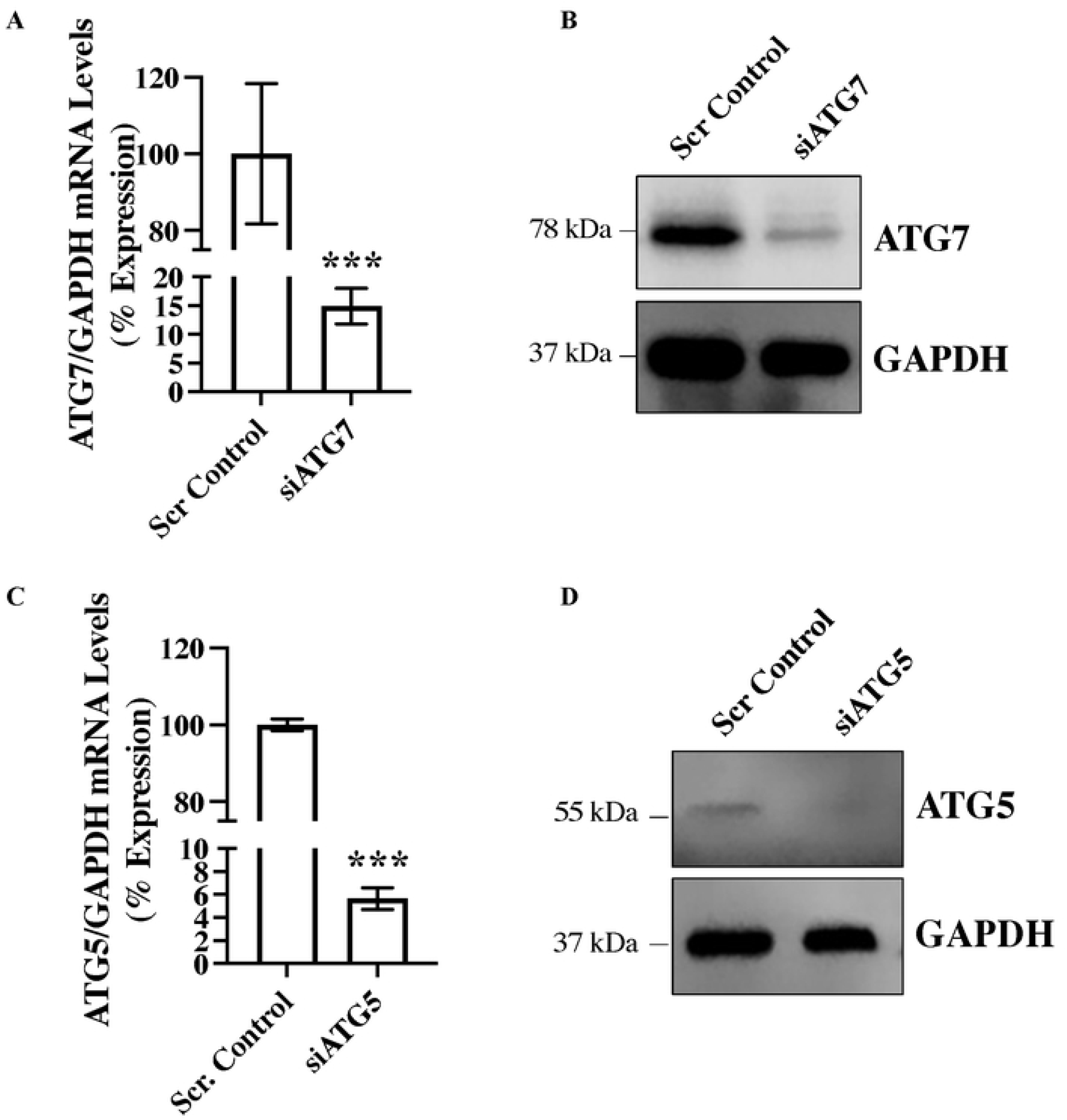
Successful silencing of ATG7 and ATG5 in endothelial cells. Human umbilical vein endothelial cells (HUVECs) were reverse-transfected with siRNAs targeting ATG7 (siATG7), ATG5 (siATG5), or a scrambled control. RNA and protein were collected 24 and 48 hours post-transfection, respectively, to perform RT-qPCR and western blot analyses. GAPDH was used as a housekeeping gene for qPCR and as a loading control for western blotting. **(A)** qPCR and **(B)** western blot results for ATG7, while **(C)** and **(D)** show the corresponding data for ATG5. Data represent the mean ± SD from three independent experiments conducted in triplicate (n = 3). Statistical analysis was performed using Student’s t-test, with *** indicating p < 0.001 compared to the scrambled control.

### Differential Gene Expression Analysis of the TGF-β Pathway Following ATG7 and ATG5 Silencing

To investigate the downstream effects of ATG7 and ATG5 silencing on the TGF-β signaling pathway, a focused PCR array profiling 84 TGF-β pathway-associated genes was conducted (Supplementary Table 1). In endothelial cells transfected with siRNA targeting ATG7, a total of 16 differentially expressed genes (DEGs) were identified (16/84; 19%). Of these, 12 genes were significantly upregulated (12/84; 14.2%, Table 1)[8] and 4 genes were significantly downregulated (4/84; 4.76%, Table 2). Among these, *bone morphogenetic protein receptor, type IB* (*BMPR1B)* exhibited the highest upregulation (3.84-fold), while *V-myc myelocytomatosis viral oncogene homolog* (*MYC)* was the most downregulated gene (−2.72-fold) relative to scrambled control-transfected endothelial cells. In contrast, siRNA-mediated silencing of ATG5 resulted in a more pronounced transcriptional response, with 39 genes significantly differentially expressed gene (39/84; 46.4%). This included 34 upregulated genes (34/84; 40.4%, Table 3) and 5 downregulated genes (5/84; 5.95%, Table 4). The most upregulated gene was *Inhibin, alpha* (*INHA*; 7.42-fold), while *TSC22 domain family, member 1* (*TSC22D1)* was the most downregulated (−4.46-fold) in siATG5-transfected cells compared to scrambled controls.

**Table 1.**
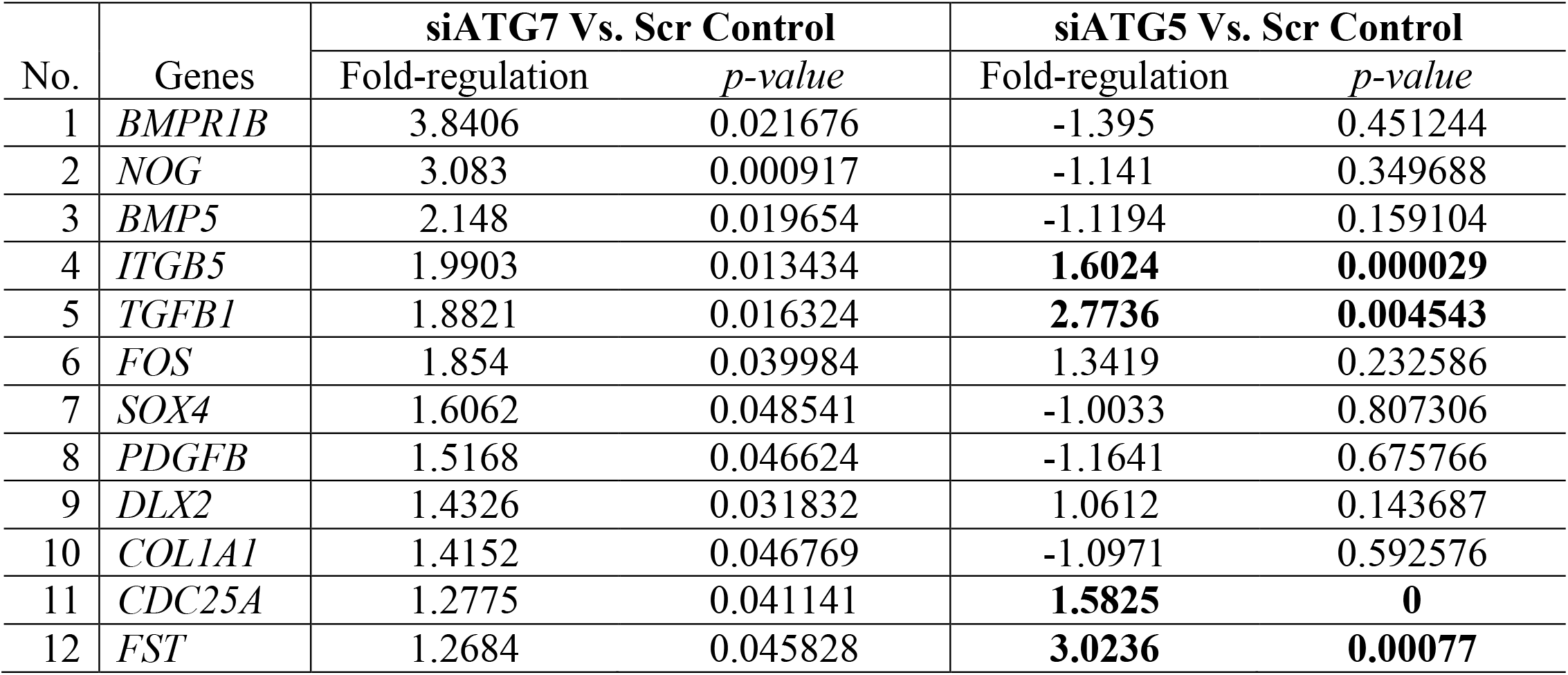
Genes up-regulated in ATG7-silenced vs. control endothelial cells and their status in ATG5-silenced endothelial cells. The data was collected 24 hours post-transfection using the TGF-ß/BMP signalling pathway PCR array. *Bone Morphogenetic Protein Receptor Type 1B (BMPR1B); Noggin (NOG); Bone Morphogenetic Protein 5 (BMP5); Integrin Subunit Beta 5 (ITGB5); Transforming Growth Factor Beta 1 (TGFB1); Fos Proto-Oncogene, AP-1 Transcription Factor Subunit (FOS); SRY-Box Transcription Factor 4 (SOX4); Platelet Derived Growth Factor Subunit B (PDGFB); Distal-Less Homeobox 2 (DLX2); Collagen Type I Alpha 1 Chain (COL1A1); Cell Division Cycle 25A (CDC25A); and Follistatin (FST)*.

**Table 2.**
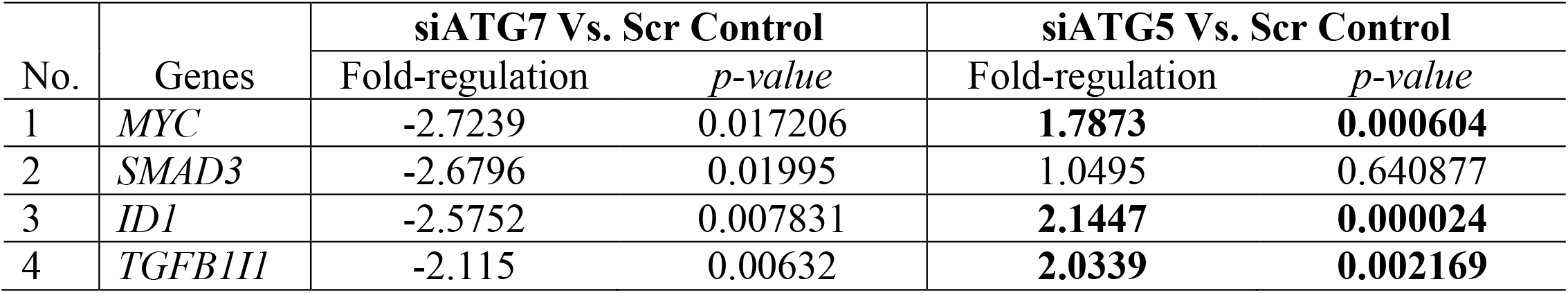
Genes down-regulated in ATG7-silenced vs. control endothelial cells and their status in ATG5-silenced endothelial cells. The data was collected 24 hours post-transfection using the TGF-ß/BMP signalling pathway PCR array. *MYC Proto-Oncogene, BHLH Transcription Factor (MYC); SMAD Family Member 3 (SMAD3); Inhibitor of DNA Binding 1, HLH Protein (ID1); and Transforming Growth Factor Beta 1 Induced Transcript 1 (TGFB1I1)*.

**Table 3.**
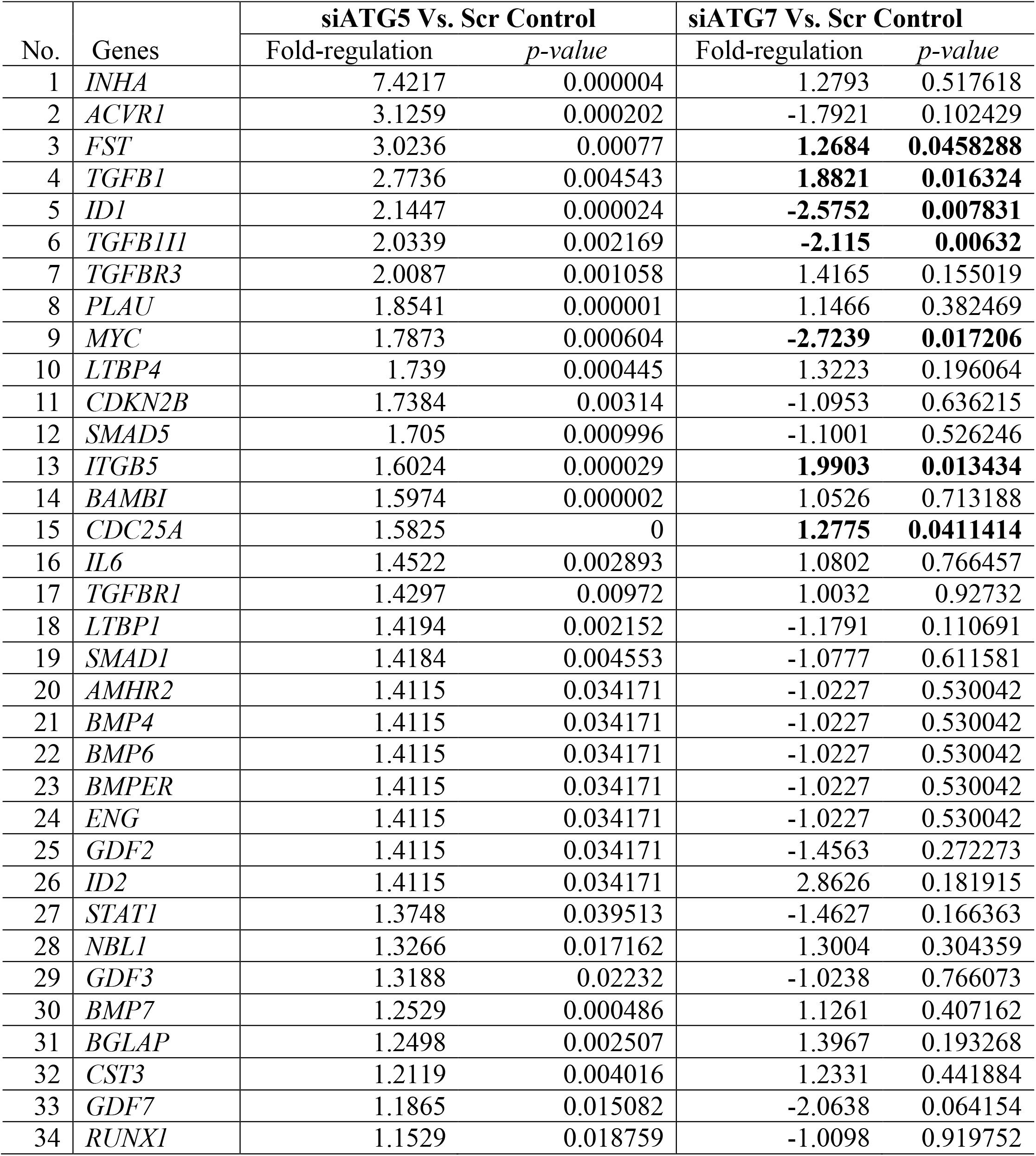
Genes up-regulated in ATG5-silenced vs. control endothelial cells and their status in ATG7-silenced endothelial cells. The data was collected 24 hours post-transfection using the TGF-ß/BMP signalling pathway PCR array. *Inhibin subunit alpha (INHA); Activin A receptor type I (ACVR1); Follistatin (FST); Transforming growth factor beta 1 (TGFB1); Inhibitor of DNA binding 1, HLH protein (ID1); Transforming growth factor beta 1 induced transcript 1 (TGFB1I1); Transforming growth factor beta receptor III (TGFBR3); Plasminogen activator, urokinase (PLAU); MYC proto-oncogene, BHLH transcription factor (MYC); Latent transforming growth factor beta binding protein 4 (LTBP4); Cyclin dependent kinase inhibitor 2B (CDKN2B); SMAD family member 5 (SMAD5); Integrin subunit beta 5 (ITGB5); BMP and activin membrane-bound inhibitor homolog (BAMBI); Cell division cycle 25A (CDC25A); Interleukin 6 (IL6); Transforming growth factor beta receptor I (TGFBR1); Latent transforming growth factor beta binding protein 1 (LTBP1); SMAD family member 1 (SMAD1); Anti-Müllerian hormone receptor type 2 (AMHR2); Bone morphogenetic protein 4 (BMP4); Bone morphogenetic protein 6 (BMP6); BMP-binding endothelial regulator (BMPER); Endoglin (ENG); Growth differentiation factor 2 (GDF2); Inhibitor of DNA binding 2, HLH protein (ID2); Signal transducer and activator of transcription 1 (STAT1); Neuroblastoma suppression of tumorigenicity 1 (DAN family BMP antagonist) (NBL1); Growth differentiation factor 3 (GDF3); Bone morphogenetic protein 7 (BMP7); Bone gamma-carboxyglutamate protein (osteocalcin) (BGLAP); Cystatin C (CST3); Growth differentiation factor 7 (GDF7); and Runt related transcription factor 1 (RUNX1)*.

**Table 4.** Genes down-regulated in ATG5-silenced vs. control endothelial cells and their status in ATG7-silenced endothelial cells. The data was collected 24 hours post-transfection using the TGF-ß/BMP signalling pathway PCR array. *TSC22 Domain Family Member 1 (TSC22D1); SMAD Family Member 4 (SMAD4); Homeodomain Interacting Protein Kinase 2 (HIPK2); Growth Differentiation Factor 5 (GDF5); and Collagen Type III Alpha 1 Chain (COL3A1)*.

| No. | Genes | siATG5 Vs. Scr Control |  | siATG7 Vs. Scr Control |  |
| --- | --- | --- | --- | --- | --- |
|  |  | Fold-regulation | *p-value* | Fold-regulation | *p-value* |
| 1 | *TSC22D1* | -4.4637 | 0.017904 | 1.4533 | 0.373053 |
| 2 | *SMAD4* | -3.1728 | 0.046026 | 1.4185 | 0.385824 |
| 3 | *HIPK2* | -1.3902 | 0.005432 | -1.8455 | 0.147838 |
| 4 | *GDF5* | -1.2451 | 0.0056 | 1.2241 | 0.260439 |
| 5 | *COL3A1* | -1.2382 | 0.041741 | -1.4009 | 0.10176 |

Among the 12 upregulated DEGs identified in siATG7-transfected endothelial cells, four genes— *integrin, beta 5* (*ITGB5*; 1.99-fold), *transforming growth factor, beta 1* (*TGFB1*; 1.88-fold), *cell division cycle 25 homolog A* (*CDC25A*; 1.27-fold), and *follistatin* (*FST*; 1.26-fold)—were also found to be upregulated in siATG5-transfected cells, with fold changes of 1.60, 2.70, 1.58, and 3.00, respectively (Table 1). Interestingly, among the four downregulated DEGs in the siATG7-transfected condition, three genes—*MYC* (−2.72-fold), *inhibitor of DNA binding 1, dominant negative helix-loop-helix protein* (*ID1*; −2.14-fold), *and transforming growth factor beta 1 induced transcript 1* (*TGFB1I1*; −2.11-fold)—exhibited significant upregulation in siATG5-transfected endothelial cells, with respective fold changes of 1.78, 2.14, and 2.03 (Table 2). Among the 34 significantly upregulated DEGs identified in siATG5-transfected endothelial cells, seven genes were also differentially expressed in siATG7-transfected cells. Of these, four genes—*FST* (3.02-fold), *TGFB1* (2.77-fold), *ITGB5* (1.60-fold), and *CDC25A* (1.58-fold)—were likewise upregulated in siATG7-transfected cells, with corresponding fold changes of 1.26, 1.88, 1.99, and 1.27, respectively (Table 3). Given the important role of TGFB1 in EndMT and endothelial function, we validated our PCR array data, which also confirmed TGFB1 upregulation in siATG7-transfected (fold change 1.51±0.25, *p* = 0.02 *versus* scrambled control) and siATG5-transfected (fold change 1.57±0.13, *p* = 0.0001 versus scrambled control) endothelial cells. In contrast, the remaining three genes—*ID1* (2.14-fold), *TGFB1I1* (2.03-fold), and MYC (1.78-fold)—which were upregulated in the siATG5 condition, were significantly downregulated in siATG7-transfected endothelial cells, with fold changes of −2.14, −2.11, and −2.72, respectively. This divergence highlights distinct regulatory consequences of ATG5 and ATG7 silencing on specific TGF-β pathway genes. Notably, among the five significantly downregulated DEGs identified in siATG5-transfected endothelial cells, none were found to be significantly differentially expressed in siATG7-transfected cells (Table 4), suggesting that these gene expression changes may be uniquely associated with ATG5 silencing.

### ATG7 and ATG5 Differentially Regulate Endothelial Function

Autophagy is a key regulator of endothelial cell function, modulating processes such as proliferation and migration through its roles in energy homeostasis, protein turnover, and intracellular signaling pathways [16-18]. Accordingly, assays were performed to evaluate the effect of ATG7 and ATG5 loss on endothelial cell migration and proliferation. Endothelial cell-specific loss of ATG7 has been shown to induce TGF-β-driven EndMT [8]. Moreover, TGF-β-induced EndMT has been reported to impair endothelial proliferation and enhance migratory activity [15, 16]. Surprisingly, however, ATG7-deficient endothelial cells exhibited a reduced migratory capacity (Fig. 2A and B). Interestingly, loss of ATG5 resulted in a significantly increased migratory capacity, both compared to scrambled control and to ATG7-deficient endothelial cells (Fig. 2A and B). As expected, loss of ATG7 was associated with a reduction in proliferative capacity. While ATG5-deficient endothelial cells showed significantly higher proliferation than ATG7-deficient cells, their proliferation was comparable to that of control cells (Fig. 3A). Therefore, loss of ATG5 does not appear to have a significant effect on endothelial cell proliferation. The cyclin-dependent kinase inhibitor p21 (Cip1/Waf1) restrains endothelial cell proliferation by inhibiting CDK–cyclin complexes and interfering with PCNA binding, thereby enforcing G1 phase cell cycle arrest [19]. To investigate whether p21 is differentially regulated by ATG7 and ATG5, we performed qPCR and Western blot analysis in scrambled control, siATG7-, and siATG5-transfected endothelial cells. In line with the reduced proliferation observed in ATG7-deficient and unaffected proliferation in ATG5-deficient endothelial cells, p21 expression was significantly increased at both the transcript and protein levels in ATG7-deficient cells, while no change was observed in ATG5-deficient cells (Fig. 3B, C and D).

**Figure 2.**
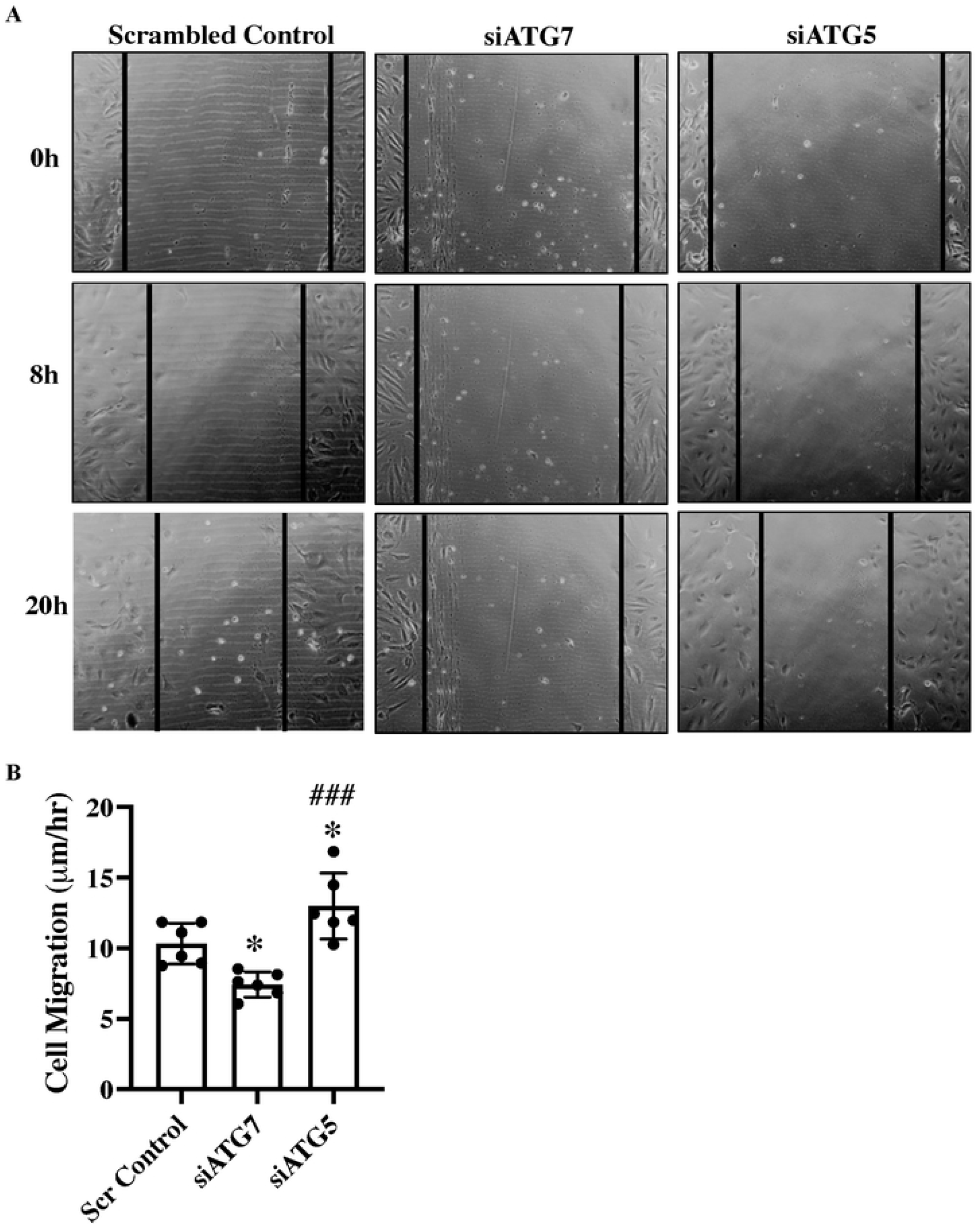
Differences in migration abilities of HUVECs following silencing of ATG7 and ATG5. **(A)** A scratch wound was created in the cell monolayer of scrambled control, siATG7, and siATG5-transfected HUVECs using a p200 pipette tip. Cells were incubated in low serum media (MCDB131 + 1% FBS) and imaged immediately at time zero (T0) and every 4 hours up to 20 hours using phase contrast microscopy. The percentage of open wound area at each time point was quantified using the ImageJ wound healing tool. **(B)** Migration velocity, defined as the rate at which cells closed the scratch wound, was calculated for each group. Data are expressed as mean ± SD and analyzed by one-way ANOVA with Tukey’s multiple comparison test. Statistical significance is indicated as *p < 0.05 *versus* scrambled control, and ###p < 0.001 *versus* siATG7.

**Figure 3.**
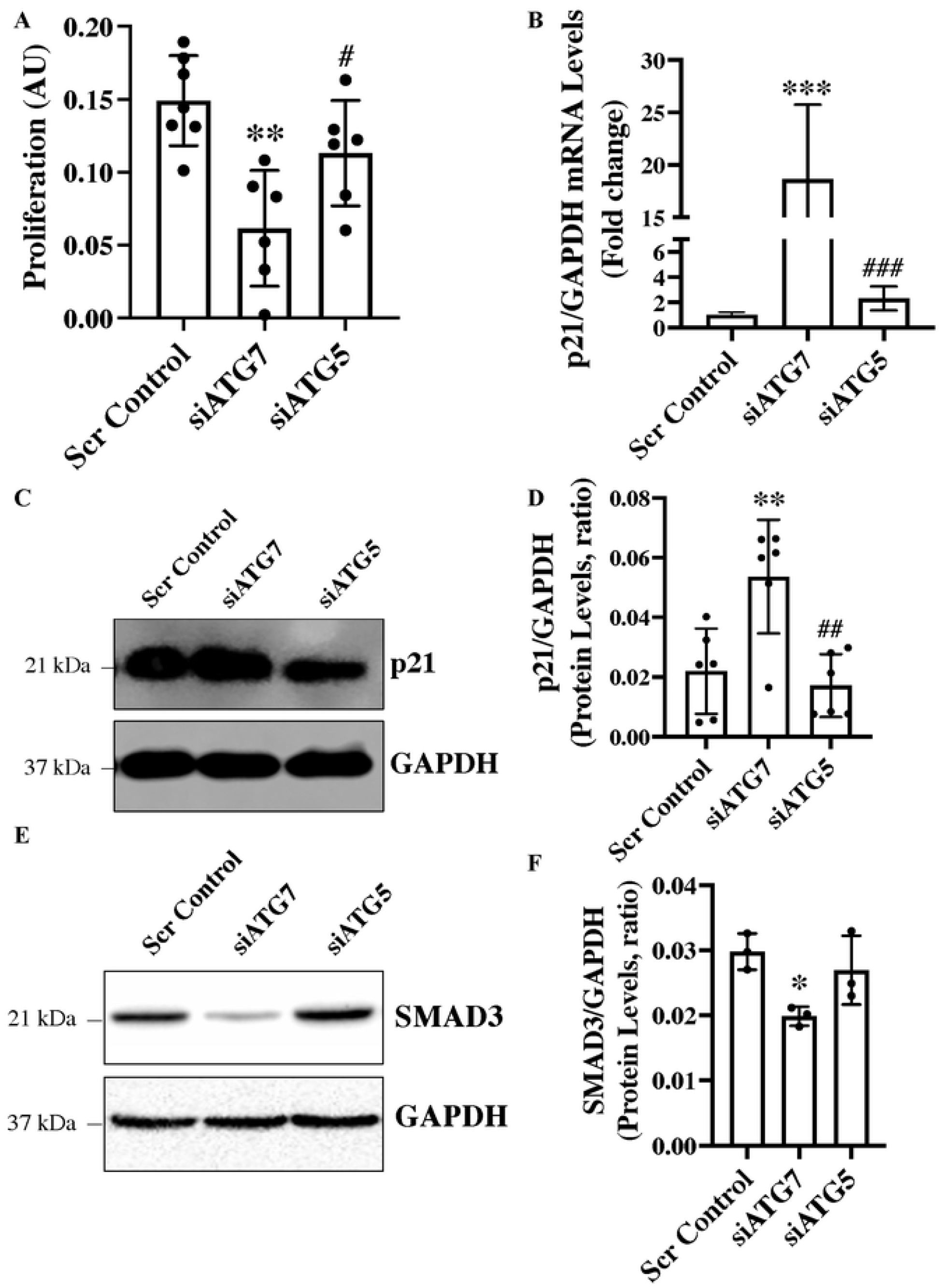
Differences in proliferation and expression of the cell cycle inhibitor p21 in HUVECs following silencing of ATG7 and ATG5. **(A)** HUVECs were transfected with siATG7, siATG5, or scrambled control in 96-well plates for 72 hours, followed by addition of WST-1 reagent. After 4 hours incubation at 37°C with 5% CO2, absorbance was measured at 440 nm with 655 nm as a reference wavelength to assess cell proliferation. For gene and protein expression analyses, HUVECs were reverse transfected with siRNAs, with RNA collected 24 hours and protein collected 48 hours post-transfection. **(B)** RT-qPCR, **(C)** western blot, and **(D)** quantification analyses were performed for p21 expression. **(E)** Western blot and **(F)** quantification were also conducted for SMAD3. GAPDH was used as a loading control. Data are presented as mean ± SD and analyzed by one-way ANOVA with Tukey’s multiple comparison test. Statistical significance is indicated by *p < 0.05, **p < 0.01 and ***p < 0.001 *versus* scrambled control, and #p < 0.05, ##p < 0.01 and ###p < 0.001 *versus* siATG7.

## Discussion

This study reveals that although ATG5 and ATG7 are both indispensable for canonical autophagy, their loss in endothelial cells elicits markedly distinct downstream effects, particularly in the context of TGF-β signaling and endothelial function. Consistent with their shared role in autophagosome formation, silencing of either gene disrupted autophagy; however, only ATG7 deficiency induced transcriptional reprogramming characteristic of EndMT, along with impaired endothelial proliferation and reduced migration. These findings reinforce the idea that ATG7 plays a unique and non-redundant role in maintaining endothelial phenotype and function, potentially through autophagy-independent mechanisms. In contrast, ATG5 silencing, while als autophagy, did not promote EndMT or impair proliferation—instead, it led to a significant increase in endothelial cell migration without affecting proliferation. The TGF-β PCR array was essential to determine whether loss of ATG7 and ATG5, despite both blocking autophagy, would differentially affect the transcriptional landscape of the TGF-β signaling pathway, which is closely linked to endothelial dysfunction and EndMT. This approach allowed us to uncover distinct gene expression profiles and regulatory outcomes associated with each gene’s silencing, highlighting potential autophagy-independent roles of ATG7 and ATG5 in modulating TGF-β-driven responses in endothelial cells. Specifically, ATG7 deficiency resulted in a modest alteration of gene expression, with only 12 out of 84 TGF-β pathway-associated genes (19%) being differentially expressed. In contrast, ATG5 silencing triggered a substantially broader transcriptional shift, with 39 genes (46.4%) significantly affected—more than three times the number observed in ATG7-deficient cells. Notably, the majority of these DEGs in both conditions were upregulated, yet the magnitude and diversity of the response were much greater in ATG5-deficient cells, where 34 genes were upregulated versus only 12 in the ATG7-deficient condition. This broader response in the absence of ATG5 may reflect distinct compensatory or regulatory mechanisms that are activated upon ATG5 loss, potentially indicating roles for ATG5 beyond canonical autophagy. Conversely, the more restricted DEG profile in ATG7-deficient cells, despite its critical upstream position in the autophagy cascade, suggests that ATG7 may exert more targeted or context-specific regulatory effects, especially in relation to TGF-β signaling. These findings support the idea that ATG5 and ATG7 have non-overlapping functions, and that the extent of gene expression remodeling in response to their loss is not merely a reflection of autophagy impairment, but may also involve autophagy-independent roles unique to each protein.

The gene expression profile observed in ATG7-deficient endothelial cells suggests a pro-EndMT and anti-proliferative, anti-migratory phenotype, consistent with the functional impairments identified in these cells. Several upregulated genes are closely associated with mesenchymal activation and EndMT. For example, BMPR1B, BMP5, and TGFB1 are ligands and receptors within the TGF-β/BMP signaling axis, known to drive EndMT by activating SMAD-dependent and SMAD-independent pathways that repress endothelial markers and promote mesenchymal gene expression [20, 21]. SOX4 and FOS, transcription factors upregulated in this context, have been implicated in promoting mesenchymal transition and fibrosis in various endothelial and epithelial systems [22, 23]. Moreover, PDGFB and COL1A1, both elevated in ATG7-deficient cells, are canonical mesenchymal markers often associated with extracellular matrix deposition and pericyte-like phenotypes [24]. NOG and FST, which encode extracellular inhibitors of BMPs, may further disrupt the finely tuned balance of pro- and anti-EndMT signaling, creating a permissive environment for mesenchymal transition [25]. Concurrently, the downregulation of MYC, a key driver of cell proliferation, aligns with the observed reduction in endothelial growth in ATG7-deficient cells. MYC represses cell cycle inhibitors like p21 and promotes endothelial proliferation and angiogenesis; thus, its suppression may contribute directly to the growth arrest phenotype [26, 27]. We indeed observed reduced p21 expression and increased proliferation in ATG7-deficient endothelial cells. Additionally, reduced expression of ID1, a downstream target of BMP signaling that normally maintains endothelial identity and promotes proliferation and migration, has been shown to facilitate EndMT and inhibit endothelial cell proliferation [28]. SMAD3 downregulation in this context may appear paradoxical, as SMAD3 typically promotes EndMT. Accordingly, we confirmed whether the reduced SMAD3 transcription leads to reduced SMAD3 protein, we performed western blot for SMAD3, which confirmed SMAD3 reduction at protein levels (Fig. 3E and F). However, its suppression may reflect a shift toward non-canonical TGF-β signaling or feedback inhibition mechanisms [29]. Finally, TGFB1I1, also known as Hic-5, is a LIM domain protein that regulates focal adhesions and cytoskeletal remodeling; its downregulation may contribute to the reduced migratory capacity observed in ATG7-deficient cells [30]. Collectively, these transcriptional changes suggest that loss of ATG7 leads to a gene expression program favoring mesenchymal transition while impairing cell proliferation and migration.

The transcriptomic profile of ATG5-deficient endothelial cells reveals a gene expression pattern that, despite extensive remodeling of the TGF-β signaling pathway, does not favor EndMT. Several key pro-EndMT regulators such as SMAD3, ZEB1, or SNAIL were not upregulated, and notably, SMAD4—a central mediator of canonical TGF-β signaling—was significantly downregulated. This likely disrupted the downstream transcriptional programs necessary for initiating mesenchymal transition [31]. Furthermore, the upregulation of BMP pathway components like BMP4, BMP6, BMP7, SMAD1, and SMAD5, alongside BMP antagonists such as FST, NBL1, and BAMBI, suggests a shift toward BMP-dominant signaling, which has been shown to counteract TGF-β-induced EndMT in endothelial cells [20, 32]. This BMP-skewed balance, together with preserved or upregulated expression of anti-EndMT regulators such as ID1, ID2, and MYC—all of which maintain endothelial identity and suppress mesenchymal reprogramming [28, 33]—likely explains why ATG5-deficient cells fail to undergo EndMT despite upregulation of TGFB1 and its receptors.

Interestingly, ATG5-deficient cells displayed enhanced migratory capacity, which is supported by the upregulation of genes involved in cytoskeletal remodeling, matrix degradation, and motility. For example, PLAU promotes pericellular matrix degradation, facilitating cell movement [34], while ITGB5 (integrin β5) supports dynamic interactions with the extracellular matrix critical for migration [35]. Upregulation of TGFB1I1, a focal adhesion adaptor, further supports enhanced migration through its role in regulating actin stress fibers and focal adhesion turnover [30]. Additionally, ENG, although classically involved in vascular development, can modulate TGF-β/BMP balance and enhance migratory responses under certain contexts [36]. The concurrent upregulation of pro-inflammatory mediators like IL6 and transcription factors such as STAT1 may also contribute to enhanced migratory signaling networks. Despite these broad changes, cell proliferation remained unaffected in ATG5-deficient cells, likely due to a balanced regulation of cell cycle genes. While pro-proliferative genes such as MYC and CDC25A were upregulated, the expression of cell-cycle inhibitors like CDKN2B was also increased. This suggests a compensatory mechanism, where proliferative signals are counteracted by inhibitory cues, resulting in a net neutral effect on endothelial proliferation [37]. Additionally, p21—a key regulator that induces cell cycle arrest—was upregulated in ATG7-deficient cells but remained unchanged in ATG5-deficient cells, which is consistent with the normal proliferation observed in ATG5-deficient endothelial cells.

The distinct transcriptional landscapes observed in ATG7- and ATG5-deficient endothelial cells highlight their non-redundant roles in modulating TGF-β/BMP signaling and endothelial phenotype. While both ATG proteins are essential for canonical autophagy, the silencing of ATG7 led to selective yet profound changes in genes associated with EndMT, proliferation, and migration. Conversely, ATG5-deficient endothelial cells exhibited a broader and more heterogeneous transcriptional response, with upregulation of pro-migratory and pro-survival genes, which may underlie the enhanced migratory phenotype observed. Taken together, these findings suggest that ATG7 and ATG5 differentially regulate the TGF-β/BMP axis, possibly through both autophagy-dependent and independent mechanisms. The selective upregulation or downregulation of overlapping genes—such as TGFB1, FST, ITGB5, and CDC25A, which are upregulated in both knockdowns—may reflect shared responses to autophagy disruption. However, the opposing regulation of key transcriptional regulators and signaling mediators such as MYC, ID1, and TGFB1I1 likely drives the divergent phenotypic outcomes. Specifically, ATG7 loss promotes EndMT, suppresses proliferation, and reduces migration, while ATG5 loss enhances migration without triggering EndMT or impairing proliferation. These results underscore the complex and gene-specific regulatory roles of autophagy proteins in endothelial cell biology and vascular remodeling, with potential implications for targeting autophagy pathways in vascular disease.

In summary, this study reveals that while both ATG5 and ATG7 are essential for autophagy, their loss leads to distinct effects on endothelial cell function and TGF-β/BMP signaling. ATG7 deficiency promotes EndMT and impairs proliferation and migration, whereas ATG5 loss enhances migration without inducing EndMT. These findings highlight gene-specific, potentially autophagy-independent roles of ATG5 and ATG7 in regulating endothelial phenotype. Although human umbilical vein endothelial cells (HUVECs) represent a well-established and widely utilized in vitro model for studying endothelial biology, they capture only a subset of the phenotypic and molecular diversity inherent to the endothelium across different vascular beds. Endothelial heterogeneity is well-documented, with cells from arterial, venous, microvascular, and organ-specific regions exhibiting distinct gene expression profiles, signaling responses, and physiological roles [38, 39]. As such, findings derived from HUVECs require further validation in endothelial cells from other vascular territories, including aortic, microvascular, and macrovascular origins, to ensure broader biological relevance. Furthermore, while this study provides valuable insights into transcriptional changes under the experimental conditions, mRNA expression levels do not always correspond to protein abundance or functional activity due to the influence of post-transcriptional, translational, and post-translational regulatory mechanisms [40]. Therefore, further validation at the protein level is essential to confirm the biological significance of the DEGs. Additionally, to determine whether the observed changes are specifically attributable to the loss of ATG5 or ATG7, these transcriptional alterations should be validated using pharmacological modulators of autophagy in endothelial cells. This would help distinguish gene expression changes that are autophagy-dependent from those that may be secondary or compensatory effects.

**Supplementary Table 1**. List of 84 genes present in TGF-ß/BMP signalling pathway PCR array.

## References

1. Abbas, T. and A. Dutta, p21 in cancer: intricate networks and multiple activities. Nat Rev Cancer, 2009. 9(6): p. 400–14.

2. Furuya, K., et al., Effects of GDF7/BMP12 on proliferation and alkaline phosphatase expression in rat osteoblastic osteosarcoma ROS 17/2.8 cells. J Cell Biochem, 1999. 72(2): p. 177–80.

3. Mizushima, N., et al., Dissection of autophagosome formation using Apg5-deficient mouse embryonic stem cells. J Cell Biol, 2001. 152(4): p. 657–68.

4. Tanida, I., T. Ueno, and E. Kominami, Human light chain 3/MAP1LC3B is cleaved at its carboxyl-terminal Met121 to expose Gly120 for lipidation and targeting to autophagosomal membranes. J Biol Chem, 2004. 279(46): p. 47704–10.

5. Fujita, N., et al., The Atg16L complex specifies the site of LC3 lipidation for membrane biogenesis in autophagy. Mol Biol Cell, 2008. 19(5): p. 2092–100.

6. Mizushima, N., et al., Mouse Apg16L, a novel WD-repeat protein, targets to the autophagic isolation membrane with the Apg12-Apg5 conjugate. J Cell Sci, 2003. 116(Pt 9): p. 1679–88.

7. Komatsu, M., et al., Impairment of starvation-induced and constitutive autophagy in Atg7-deficient mice. J Cell Biol, 2005. 169(3): p. 425–34.

8. Singh, K.K., et al., The essential autophagy gene ATG7 modulates organ fibrosis via regulation of endothelial-to-mesenchymal transition. J Biol Chem, 2015. 290(5): p. 2547–59.

9. Torisu, T., et al., Autophagy regulates endothelial cell processing, maturation and secretion of von Willebrand factor. Nat Med, 2013. 19(10): p. 1281–7.

10. Grootaert, M.O., et al., Defective autophagy in vascular smooth muscle cells accelerates senescence and promotes neointima formation and atherogenesis. Autophagy, 2015. 11(11): p. 2014–2032.

11. Nussenzweig, S.C., S. Verma, and T. Finkel, The role of autophagy in vascular biology. Circ Res, 2015. 116(3): p. 480–8.

12. Hu, M., J.M. Ladowski, and H. Xu, The Role of Autophagy in Vascular Endothelial Cell Health and Physiology. Cells, 2024. 13(10).

13. Vion, A.C., et al., Autophagy is required for endothelial cell alignment and atheroprotection under physiological blood flow. Proc Natl Acad Sci U S A, 2017. 114(41): p. E8675–E8684.

14. Torisu, K., et al., Intact endothelial autophagy is required to maintain vascular lipid homeostasis. Aging Cell, 2016. 15(1): p. 187–91.

15. Nguyen, H.C., et al., Loss of fatty acid binding protein 3 ameliorates lipopolysaccharide-induced inflammation and endothelial dysfunction. J Biol Chem, 2023. 299(3): p. 102921.

16. Singh, A., et al., Endothelial-to-Mesenchymal Transition in Cardiovascular Pathophysiology. Int J Mol Sci, 2024. 25(11).

17. Li, X., S. He, and B. Ma, Autophagy and autophagy-related proteins in cancer. Mol Cancer, 2020. 19(1): p. 12.

18. Schaaf, M.B., et al., Autophagy in endothelial cells and tumor angiogenesis. Cell Death & Differentiation, 2019. 26(4): p. 665–679.

19. Rossig, L., et al., Akt-dependent phosphorylation of p21(Cip1) regulates PCNA binding and proliferation of endothelial cells. Mol Cell Biol, 2001. 21(16): p. 5644–57.

20. Medici, D., et al., Conversion of vascular endothelial cells into multipotent stem-like cells. Nat Med, 2010. 16(12): p. 1400–6.

21. Zeisberg, M. and E.G. Neilson, Biomarkers for epithelial-mesenchymal transitions. J Clin Invest, 2009. 119(6): p. 1429–37.

22. Tiwari, N., et al., Sox4 is a master regulator of epithelial-mesenchymal transition by controlling Ezh2 expression and epigenetic reprogramming. Cancer Cell, 2013. 23(6): p. 768–83.

23. Bakiri, L., et al., Fra-1/AP-1 induces EMT in mammary epithelial cells by modulating Zeb1/2 and TGFbeta expression. Cell Death Differ, 2015. 22(2): p. 336–50.

24. Chen, P.Y., et al., Endothelial-to-mesenchymal transition drives atherosclerosis progression. J Clin Invest, 2015. 125(12): p. 4514–28.

25. Gazzerro, E. and E. Canalis, Bone morphogenetic proteins and their antagonists. Rev Endocr Metab Disord, 2006. 7(1-2): p. 51–65.

26. Dang, C.V., MYC on the path to cancer. Cell, 2012. 149(1): p. 22–35.

27. Baudino, T.A., et al., c-Myc is essential for vasculogenesis and angiogenesis during development and tumor progression. Genes Dev, 2002. 16(19): p. 2530–43.

28. Valdimarsdottir, G., et al., Stimulation of Id1 expression by bone morphogenetic protein is sufficient and necessary for bone morphogenetic protein-induced activation of endothelial cells. Circulation, 2002. 106(17): p. 2263–70.

29. Xu, J., S. Lamouille, and R. Derynck, TGF-β-induced epithelial to mesenchymal transition. Cell Research, 2009. 19(2): p. 156–172.

30. Shibanuma, M., et al., Involvement of FAK and PTP-PEST in the regulation of redox-sensitive nuclear-cytoplasmic shuttling of a LIM protein, Hic-5. Antioxid Redox Signal, 2005. 7(3-4): p. 335–47.

31. Massague, J., TGFbeta signalling in context. Nat Rev Mol Cell Biol, 2012. 13(10): p. 616–30.

32. Goumans, M.J., et al., Balancing the activation state of the endothelium via two distinct TGF-beta type I receptors. EMBO J, 2002. 21(7): p. 1743–53.

33. Kowanetz, M., et al., Id2 and Id3 define the potency of cell proliferation and differentiation responses to transforming growth factor beta and bone morphogenetic protein. Mol Cell Biol, 2004. 24(10): p. 4241–54.

34. Blasi, F. and P. Carmeliet, uPAR: a versatile signalling orchestrator. Nature Reviews Molecular Cell Biology, 2002. 3(12): p. 932–943.

35. Danen, E.H., et al., Integrins control motile strategy through a Rho-cofilin pathway. J Cell Biol, 2005. 169(3): p. 515–26.

36. Lebrin, F., et al., Endoglin promotes endothelial cell proliferation and TGF-beta/ALK1 signal transduction. EMBO J, 2004. 23(20): p. 4018–28.

37. Sherr, C.J. and J.M. Roberts, CDK inhibitors: positive and negative regulators of G1-phase progression. Genes Dev, 1999. 13(12): p. 1501–12.

38. Aird, W.C., Phenotypic heterogeneity of the endothelium: I. Structure, function, and mechanisms. Circ Res, 2007. 100(2): p. 158–73.

39. Augustin, H.G. and G.Y. Koh, Organotypic vasculature: From descriptive heterogeneity to functional pathophysiology. Science, 2017. 357(6353).

40. Liu, Y., A. Beyer, and R. Aebersold, On the Dependency of Cellular Protein Levels on mRNA Abundance. Cell, 2016. 165(3): p. 535–50.

